# Translational repression of *Nox4* mRNA via the EI24-RTRAF interaction is crucial for hydrogen peroxide homeostasis and insulin synthesis

**DOI:** 10.1101/2024.01.02.573947

**Authors:** Xintong Pei, Zhe Wang, Wenting He, Shunqin Li, Yongguang Lan, Lin Yuan, Pingyong Xu

## Abstract

As a double-edged sword, the content of reactive oxygen species (ROS) is precisely controlled. Disordered actions of ROS contribute to deleterious effects, such as cancer and metabolic dysregulation associated with aging and obesity. Although it is well established that cells have developed evolutionarily conserved programs to sense and adapt to redox fluctuations, it remains unclear how to control the expression of key ROS-producing enzymes to regulate continued ROS production at healthy levels for cells such as neurons and pancreatic beta cells. These cells have weaker antioxidant defense systems but strong secretion ability. Here, we found that the endoplasmic reticulum membrane-localized protein, EI24, controls the translation of nicotinamide adenine dinucleotide phosphate oxidase 4 (NOX4), which constitutively produces hydrogen peroxide (H_2_O_2_), by recruiting an RNA transcription, translation, and transport factor (RTRAF) to the 3’-UTRs of *Nox4*. Depletion of EI24 causes RTRAF to relocate into the nucleus, releasing the brake on *Nox4* mRNA translation, and thus, the uncontrolled translation of *Nox4* leads to a substantial generation of intracellular H_2_O_2_. This suppresses the translation of V-maf musculoaponeurotic fibrosarcoma oncogene homolog A (*MafA*), inhibits its binding to the *Ins2* gene promoter, and ultimately hinders insulin transcription. Treatment with a specific NOX4 inhibitor or the antioxidant N-acetyl-cysteine (NAC) restored *MafA* translation and downstream insulin synthesis while alleviating the diabetic symptoms in pancreatic beta-cell specific *Ei24*-KO mice. In summary, our study revealed a molecular mechanism that controls the expression of NOX4, a key enzyme responsible for continuous ROS generation. This mechanism ensures low levels of H_2_O_2_ and normal biological functions under physiological conditions.

## Introduction

Reactive oxygen species (ROS) are involved in cell signaling pathways and play crucial roles in cellular physiology and pathophysiology(Shields et al., 2021). The overproduction of ROS is associated with the oxidative damage and development of various diseases, including cancer, inflammatory diseases, and metabolic disorders associated with aging and obesity(Yang and Lian, 2020). Therefore, cells have evolved numerous antioxidant systems to maintain basal ROS at a normal, non-toxic level. This is especially essential for cells with relatively weak intrinsic antioxidant defenses, like neurons(Lee et al., 2021; Piccirillo et al., 2022) and pancreatic beta cells(Eguchi et al., 2021; Lenzen, 2017). Thus, regulating the expression levels of enzymes that constitutively produce ROS is crucial for maintaining redox homeostasis, particularly for pancreatic beta cells, that have high energy demands and engage in metabolic and redox crosstalk. The vigorous process of synthesizing and folding insulin within pancreatic beta cells results in the production of over 1 million insulin molecules per minute in a single beta cell(Scheuner and Kaufman, 2008), leading to a higher concentration of ROS produced in comparison to other cell types, as three molecules of H_2_O_2_ are generated for each insulin production. As a result, it is crucial to tightly control the generation of ROS in beta cells to ensure efficient synthesis and proper folding of insulin. However, the regulation of enzymes that constitutively produce ROS in pancreatic beta cells and its role in regulating insulin synthesis remain unknown.

Intracellular ROS mainly comprise short-lived free radicals like the superoxide radical anion (O_2•−_), hydroxyl (HO^•^) and hydroperoxyl (HO^•^_2_), as well as non-radical derivatives of O_2_ such as hydrogen peroxide (H_2_O_2_)(Shields et al., 2021). Due to its longer lifetime and the ability of freely diffuse across the cell membrane, H_2_O_2_ is an important second messenger for cell signaling given the redox status in different subcellular locations and antioxidant distribution within the cell(Pak et al., 2020; Sies, 2017). The major intracellular sources of H_2_O_2_ include mitochondrial electron transport chain, peroxisome, and the membrane-bound nicotinamide adenine dinucleotide phosphate oxidase (NOX)(Shields et al., 2021). So far, the NOX family consists of seven isoforms, with NOX4 being the only member that does not require upstream activators and is constitutively active(Wang et al., 2023). Hence, precise regulation of the expression levels NOX4 is crucial for the production H_2_O_2_. However, despite accumulating evidence that NOX4 may contribute to the occurrence and development of diseases and that its expression levels are dynamically regulated in response to a wide range of stimuli(Canugovi et al., 2019; Chen et al., 2012; Pejenaute et al., 2020; Peñuelas-Haro et al., 2023; Wang et al., 2023), the molecular mechanisms that govern NOX4 expression levels are not yet fully elucidated.

In the current study, we have discovered that the translation of *Nox4* is negatively regulated by a crucial endoplasmic reticulum (ER) membrane protein called EI24, which plays vital roles in DNA damage response and the autophagy process(Zhao et al., 2012). Previous reports have indicated a negative correlation between EI24 expression level, age, and the severity of diabetes(Yuan et al., 2018). Specifically knocking out EI24 in pancreatic beta cells leads to reduced insulin secretion, with diabetic symptoms worsening over time(Yuan et al., 2018). Here, we not only show that EI24 responds to changes in H_2_O_2_ concentration, but also demonstrate its interaction with the RNA-binding protein RTRAF to recruit it to the 3’-UTR of *Nox4* mRNA, effectively inhibiting the translation of *Nox4*. This process is critical for maintaining basal H_2_O_2_ homeostasis and insulin synthesis.

## Results

### EI24 modulates redox homeostasis of pancreatic beta cells

Transcriptome-wide analysis of islet samples from wild type (WT) and EI24 KO mice using mRNA sequencing (mRNA-seq) revealed a number of redox-related genes that were differentially expressed (*p* < 0.05), with 50 genes being upregulated and 22 genes being downregulated in *Ei24* KOs (Figure 1a, 1b and Table S1), suggesting that EI24 may be involved in regulating the redox state of pancreatic beta cells. We next measured the intracellular ROS levels of primary WT and EI24 KO pancreatic beta cells and observed a remarkable increase of ROS content in the EI24 KO group compared to WT cells. NAC, a commonly used antioxidant, was capable of decreasing ROS to extremely low levels both in WT and KO groups (Figure 1c). To confirm the effect of EI24 on the redox balance of beta cells, we generated the EI24 KO INS-1 cell lines using CRISPR- Cas9(Yuan et al., 2018) and found that loss of EI24 produced an increase in cellular ROS (Figure 1d and 1e). The excess ROS were scavenged in the KO group following treatment with NAC (Figure 1d and 1e).

**Figure 1:**
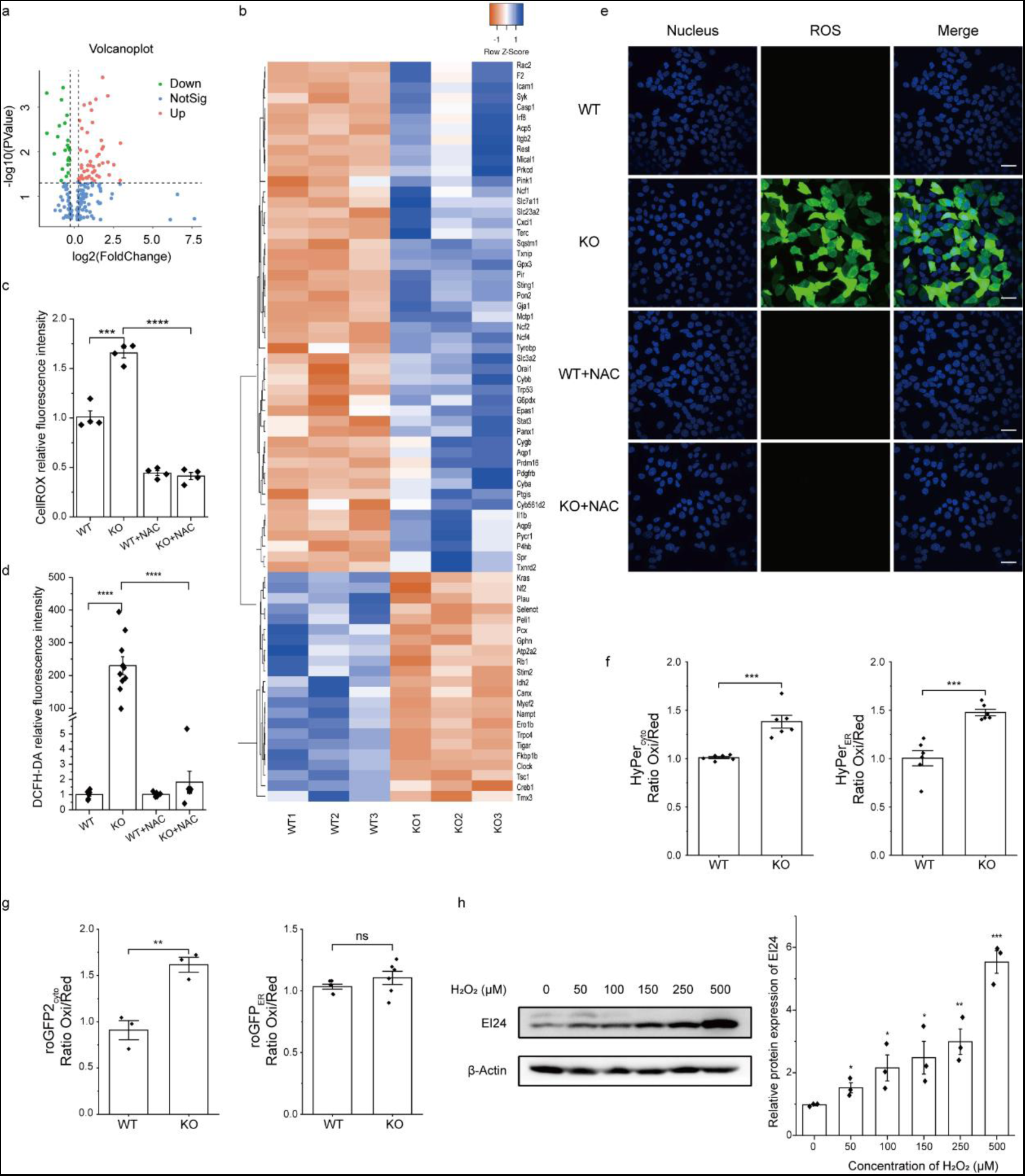
Effects of EI24 on the redox homeostasis of pancreatic beta cells. a. Scatterplot of redox-related transcripts between wild type (WT) and EI24 knockout (KO) pancreatic beta cells isolated from 12 week-old mice and sequenced with the multiplex analysis of poly(A)-linked sequences method (WT, n=3; KO, n=3). Each dot of green and red represents a gene that was significantly changed (fold change >1.5, *p* < 0.05). b. Heatmap of genes that are closely related to redox homeostasis, which are differentially upregulated or downregulated by EI24. c. Intracellular ROS production in primary pancreatic beta cells *ex vivo* with or without NAC treatment, measured as fluorescence by the CellROX kit. d. Intracellular ROS production in INS-1 cells with or without NAC treatment, measured as fluorescence intensity by the DCFH-DA probe. e. Representative images of DCFH-DA-labeled INS-1 cells. f. Relative H_2_O_2_ concentration in cytosol and ER measured with HyPer in INS-1 cells. g. Relative GSH:GSSG ratio in cytosol and ER of INS-1 cells, GSH, reduced glutathione, GSSG, and oxidized glutathione. h. EI24 levels respond to H_2_O_2_ at different concentrations. Data are expressed as the mean ± S.E.M. \**p* < 0.05, \*\**p* < 0.01, \*\*\**p* < 0.001, \*\*\*\**p* < 0.0001. Scale bars: 25 µm.

To better understand the different types and sources of excessive ROS in EI24 KO cells, and since the glutathione redox couple (GSH/GSSG) and H_2_O_2_ are central to redox homeostasis and redox signaling(Albrecht et al., 2011), the ratiometric hydrogen peroxide sensor (HyPer) and redox-sensitive green fluorescent protein (roGFP) targeted to cytosol (HyPer_cyto_/roGFP2_cyto_) and ER (HyPer_ER_/roGFP2_ER_) were used to monitor H_2_O_2_ concentration and glutathione redox state, respectively. Results showed that H_2_O_2_ concentrations were increased significantly in the cytosol and ER of the EI24 KO group compared with WT (Figure 1f). For the glutathione/glutathione disulfide (GSH/GSSG) ratios, roGFP2_cyto_ signal was elevated in the KO group, while the roGFP2_ER_ fluorescence ratio remained unaffected (Figure 1g). Interestingly, the expression levels of EI24 were positively correlated with the concentration of H_2_O_2_ in a dose-dependent manner (Figure 1h). Collectively, these results suggest that the ER membrane protein, EI24, responds to H_2_O_2_ and plays an important role in maintaining H_2_O_2_ homeostasis within the ER.

### EI24 negatively regulates NOX4 translation

The homeostasis of H_2_O_2_ is tightly regulated via processes involved in its generation and elimination. Thus, the levels of specific enzymes related to the generation of H_2_O_2_, such as Ero1α and NOX4, as well as enzymes that scavenge H_2_O_2_, including peroxiredoxins (Prxs) and glutathione peroxidases (GPxs), were assessed. We found that the expression level of NOX4 was substantially increased in KO cells (Figure 2a). The overexpression of EI24 in the KO group could decrease NOX4 expression to levels similar to that in the WT group (Figure 2b). Slight variation in the expression levels of Ero1α, Prdx4, and Gpx7/8 in KO cells was observed compared to those of the WT group (Figure S1). A selective inhibitor of NOX4, GLX351322(Anvari et al., 2015), could abolish the excess ROS in EI24 KO cells (Figure 2c, d). GLX351322 reduced H_2_O_2_ signals both in the ER and cytosol of the KO group to the levels comparable to those observed in the WT group (Figure 2e, f). These data demonstrate that EI24 knockout increases the levels of NOX4, but not that of other redox enzymes, causing the overproduction of H_2_O_2_ and the redox imbalance in both ER and cytosol.

**Figure 2:**
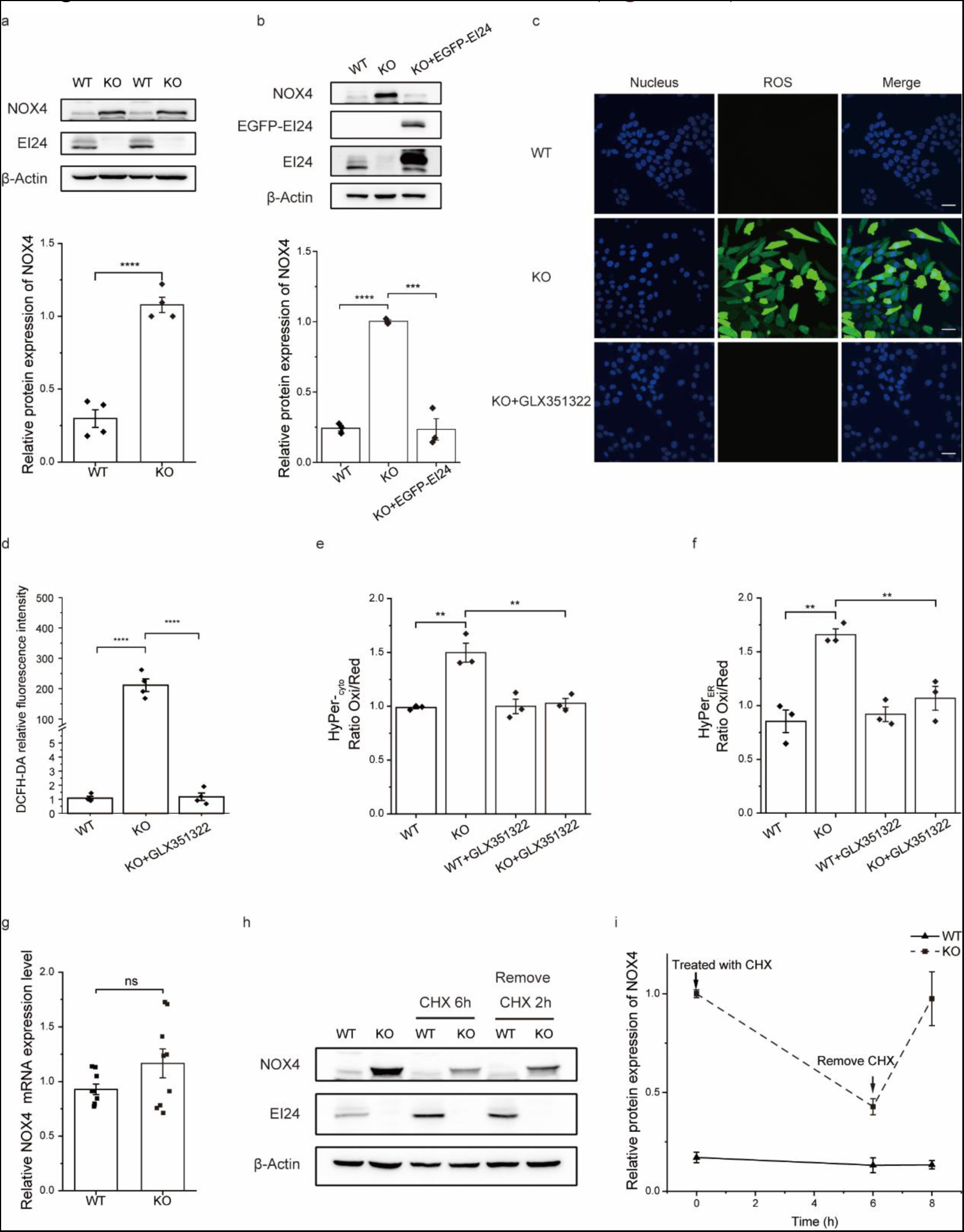
Loss of EI24 increases *Nox4* translation and H_2_O_2_ production. a. Levels of NOX4 in WT and KO INS-1 cells. b. Overexpression of EI24 in the KO group rescues NOX4 to the level of WT. c. Intracellular ROS in INS-1 cells with or without NOX4 inhibitor, GLX351322, measured as fluorescence by the DCFH-DA probe. d. Statistical analysis of fluorescence intensity of DCFH-DA probe labeled INS-1 cells with or without GLX351322 treatment. e and f. Relative H_2_O_2_ concentration in cytosol (e) and ER (f) of INS-1 cells with or without GLX351322 treatment. g. Relative *Nox4* mRNA levels in WT and EI24 KO INS-1 cells. h. Amount of NOX4 in WT and EI24 KO INS-1 cells with or without CHX treatment. i. Quantification of NOX4 from CHX chase experiments. CHX, cycloheximide. Data are expressed as the mean ± S.E.M. \*\**p* < 0.01, \*\*\**p* < 0.001, \*\*\*\**p* < 0.0001. Scale bars: 25 µm.

In order to investigate the underlying causes of heightened NOX4 expression after EI24 depletion, cells were exposed to a targeted proteasome inhibitor (MG132) as well as autophagic flux blockers (chloroquine and bafilomycin). Results showed that MG132, chloroquine, and bafilomycin had minimal impact on NOX4 expression levels (Figure S2), suggesting that the lower level of NOX4 observed in the WT group may not be attributed to protein breakdown mediated by the proteasome or autophagosomes. Interestingly, there were no noticeable differences in the mRNA levels of NOX4 between WT and KO cells (Figure 2g), indicating that EI24 may effectively regulate the translation of NOX4. This was supported by the gradual decrease in NOX4 expression over a 6-hour period when treated with a protein synthesis inhibitor, cycloheximide (CHX), in the KO group (Figure 2h). Subsequently, following the removal of CHX, the expression level of NOX4 in the KO group gradually increased. In contrast, the expression level of NOX4 in the WT group remained unchanged both during the treatment with CHX and after its removal (Figure 2h, i).

### EI24 inhibits *Nox4* translation by recruiting RTRAF to the 3’-UTRs of *Nox4*

To investigate how EI24 inhibits the translation of NOX4, we employed a dual-color fluorescence reporter system (Figure 3a). In this system, the firefly luciferase signal served as a control for normalizing expression, while the renilla luciferase functioned as a reporter in a single-read assay. As depicted in the Figure 3b, upon depletion of EI24 in cells, the introduction of the 3’-UTR motif or the combination of the 5’-UTR and 3’-UTRs of *Nox4* into the read-out system resulted in elevated ratiometric expression levels of renilla to firefly, compared to the insertion of 5’-UTR motifs alone (Figure 3b). In contrast, the control group of WT cells transfected with these three reporters individually exhibited minimal variation (Figure 3b). These findings indicate that EI24 primarily inhibits *Nox4* mRNA translation by targeting its 3’-UTRs rather than the 5’-UTRs. Additionally, we utilized the MS2-MCP system to further investigate this interaction. In this experimental system, we inserted the 3’-UTR of *Nox4* mRNA downstream of the MS2 RNA array and co-expressed it with EGFP-tagged MCP, which specifically binds MS2 RNA (Figure 3c). Through co-immunoprecipitation (IP), we successfully confirmed the interaction of EI24 with the 3’-UTR of *Nox4* mRNA (Figure 3d, e).

**Figure 3:**
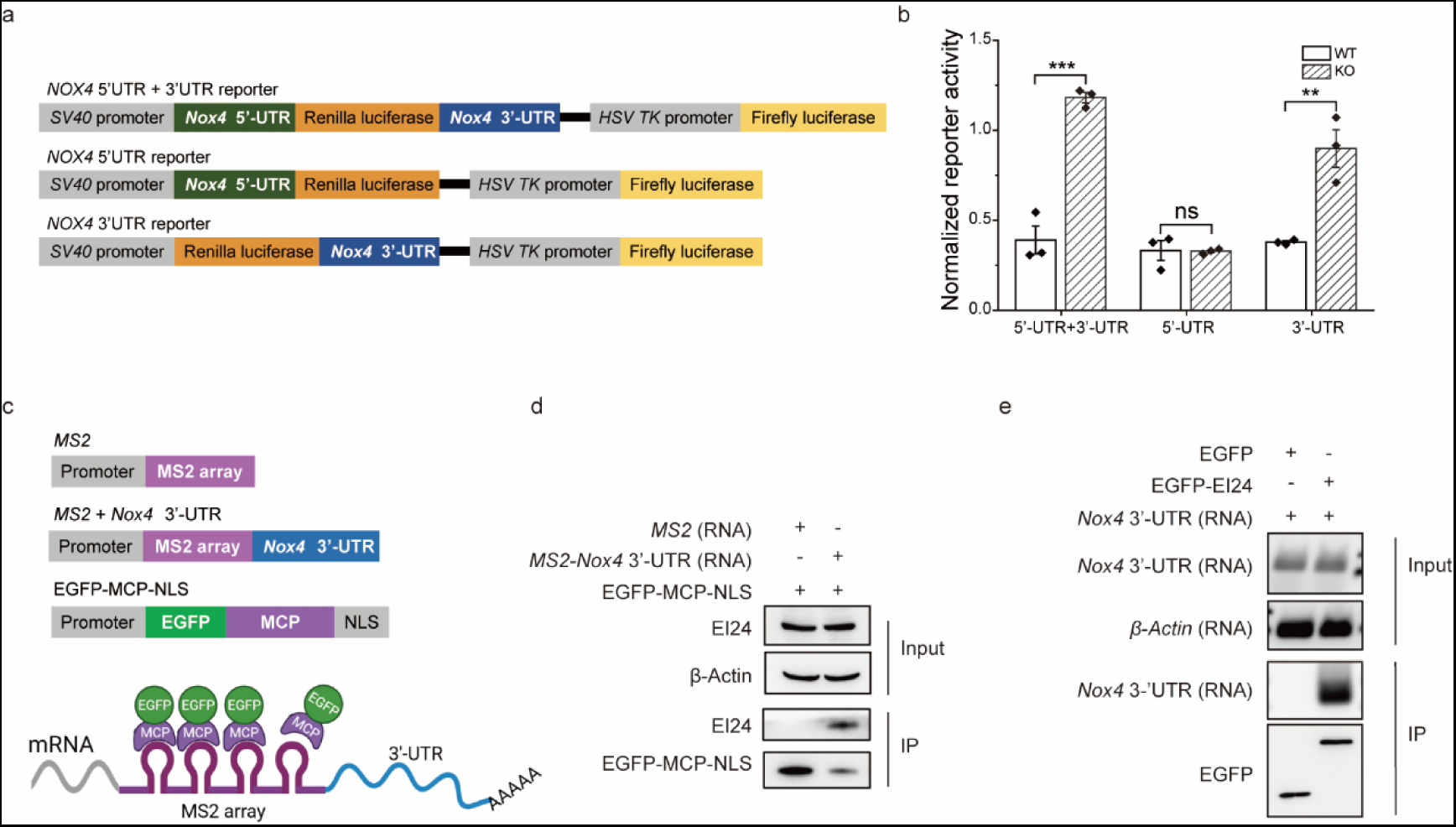
EI24 interacts with the 3’UTR of *Nox4* to inhibit its translation. a. Schematic of dual-color fluorescence reporter system. b. Relative reporter activities of the system that comprises 3’-UTR, 5’-UTR or 3’-UTR with 5’-UTR of *Nox4* mRNA in WT and KO INS-I cells. c. Schematic of MS2 assay. d and e. Co-immunoprecipitated EI24 with the 3’UTR of *Nox4* mRNA, using the 3’UTR of *Nox4* mRNA as bait (d) and EI24 as bait (e). Data are expressed as the mean ± S.E.M. \*\**p* < 0.01, \*\*\**p* < 0.001.

Because EI24 is localized in the ER membrane, we theorized that an RNA binding protein, in conjunction with EI24, could be implicated in the regulation of *Nox4* mRNA translation. Using the MS2-labeled 3’-UTR motifs of *Nox4* mRNA combined with EGFP-labeled MCP as bait, interacting proteins were detected by mass spectrometry (Figure S3). We found an RNA transcription, translation, and transport factor (RTRAF) that bound the 3’-UTRs of *Nox4* mRNA in WT cells while failing to do so in the EI24 KO group (Table S2). Furthermore, *in vitro* IP experiments also confirmed the critical role of EI24 in regulating the interaction of RTRAF with the 3’-UTR motifs of *Nox4* mRNA (Figure 4a, b). To determine whether RTRAF influences the translation of *Nox4* mRNA, we used small interfering RNA (siRNA) to knock down RTRAF in WT INS-I cells (Figure S4). Consistent with the findings observed in EI24 KO cells, RTRAF siRNA-transduced cells showed enhanced translation of the 3’-UTRs of *Nox4* mRNA (Figure 4c), accompanied by elevated NOX4 protein levels (Figure 4d), suggesting an inhibitory effect of RTRAF on *Nox4* translation. We further examined the expression level of RTRAF in EI24 KO cells and found that RTRAF was decreased by approximately 50% when EI24 is absent (Figure 4e). Interestingly, the subcellular localization of RTRAF changed after EI24 depletion, and more RTRAF protein was observed in the nucleus (Figure 4f). Nucleo-cytoplasmic partitioning also demonstrated enhanced nuclear import of RTRAF in EI24 KO cells (Figure 4g). Drawing from these observations, we hypothesize that EI24 may interact with RTRAF to trap it in the cytoplasm, mediating the binding of RTRAF to the 3’-UTR motifs of *Nox4* and inhibiting its translation. As expected, co-IP revealed that RTRAF could bind to EI24 in WT cells (Figure 4h), suggesting that EI24 and RTRAF may interact to form a protein complex. To further investigate the interaction between EI24 and RTRAF, we conducted co-IP assays to identify specific interaction domains. Based on the predicted structures of EI24 and RTRAF (UniProtKB), several truncated domains were generated, and the results in Figure 4i, 4j and supplementary figure 5 demonstrate that the N-terminal of EI24 interacts with the residues 128-178 of RTRAF (Figure 4i, 4j and Figure S5).

**Figure 4:**
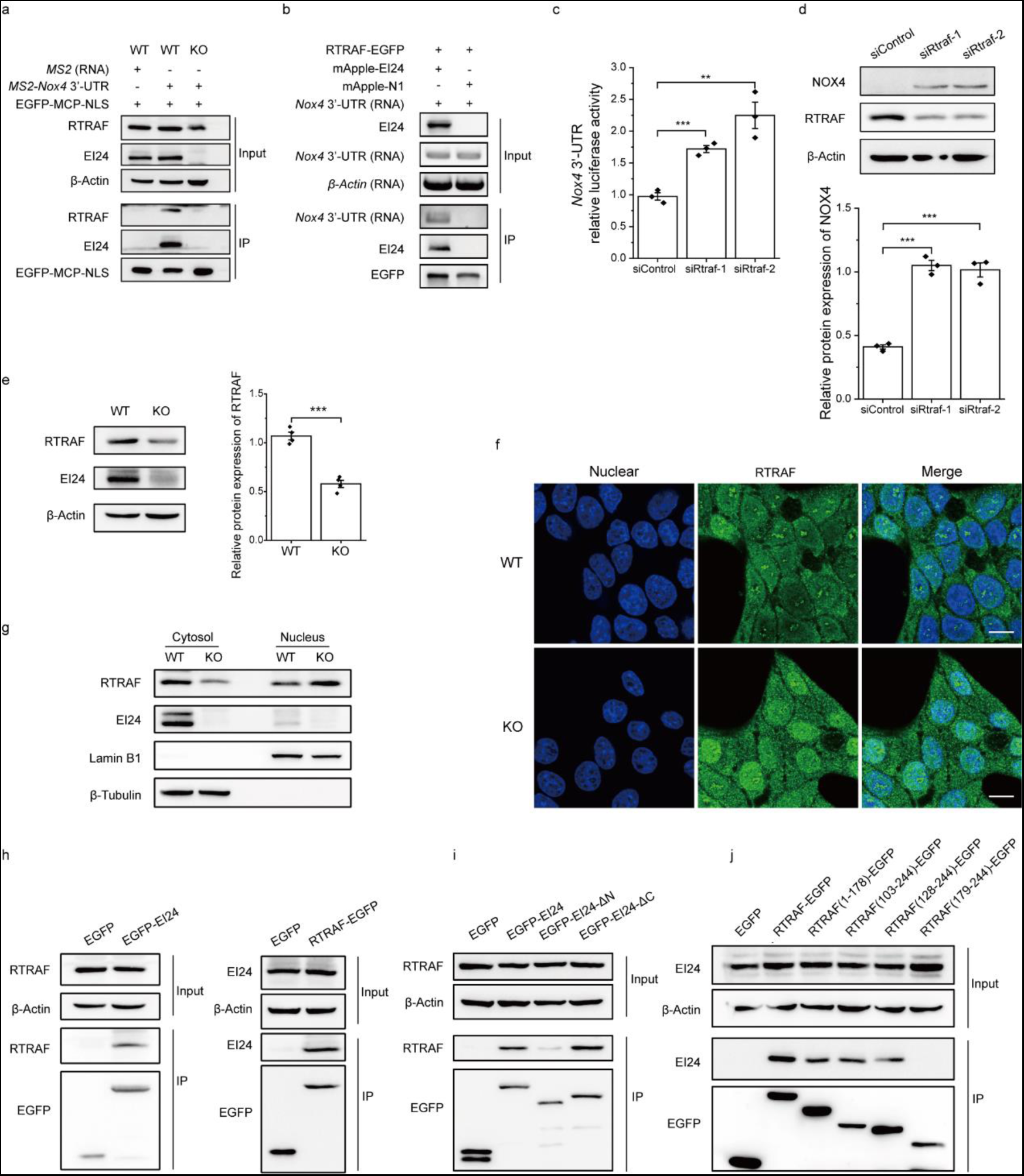
EI24 stabilizes the interaction between RTRAF and the 3’UTR of *Nox4* mRNA to inhibit *Nox4* translation. a and b. Co-immunoprecipitated RTRAF with the 3’UTR of *Nox4* mRNA in WT and EI24 KO INS-I cells, using the 3’UTR of *Nox4* mRNA as bait (a) and RTRAF as bait (b), respectively. c. Relative reporter activities of the system containing the 3’-UTR of *Nox4* mRNA in WT and EI24 KO INS-I cells with or without RTRAF siRNA treatment. d. Protein levels of NOX4 and RTRAF with or without RTRAF siRNA treatment. e. Amount of RTRAF in WT and EI24 KO INS-I cells. f. Representative images of RTRAF localization in WT and EI24 KO INS-I cells. g. Separation of nuclear and cytoplasmic proteins. h. Co-IP of EI24 with RTRAF. i and j. Identification of the interacting domains of EI24 and RTRAF by co-IP of RTRAF and EI24, using EI24 truncation mutants (i) and RTRAF truncation mutants (j), respectively. Data are expressed as the mean ± S.E.M. \*\**p* < 0.01, \*\*\**p* < 0.001, \*\*\*\**p* < 0.0001. Scale bars: 10 µm.

### EI24 knockout inhibits *Insulin* transcription and *MafA* translation by relieving *Nox4* translational inhibition

Our previous work discovered the significance of EI24 in insulin synthesis within pancreatic beta cells(Yuan et al., 2018). However, it remains unclear whether the diminished insulin synthesis in EI24 knockout cells is primarily caused by the uncontrolled translation of *Nox4*. As shown in Figure 5a, overexpression of EI24 substantially restored insulin levels in EI24 KO INS-1 cells. Conversely, expressing EGFP as a control proved ineffective in achieving the same outcome (Figure 5a). Moreover, the administration of a NOX4 inhibitor or NAC resulted in a significant recovery of insulin synthesis (Figure 5b, c). This indicates that the excessive production of H2O2, resulting from unregulated NOX4 expression following EI24 depletion, directly impacts insulin levels.

**Figure 5:**
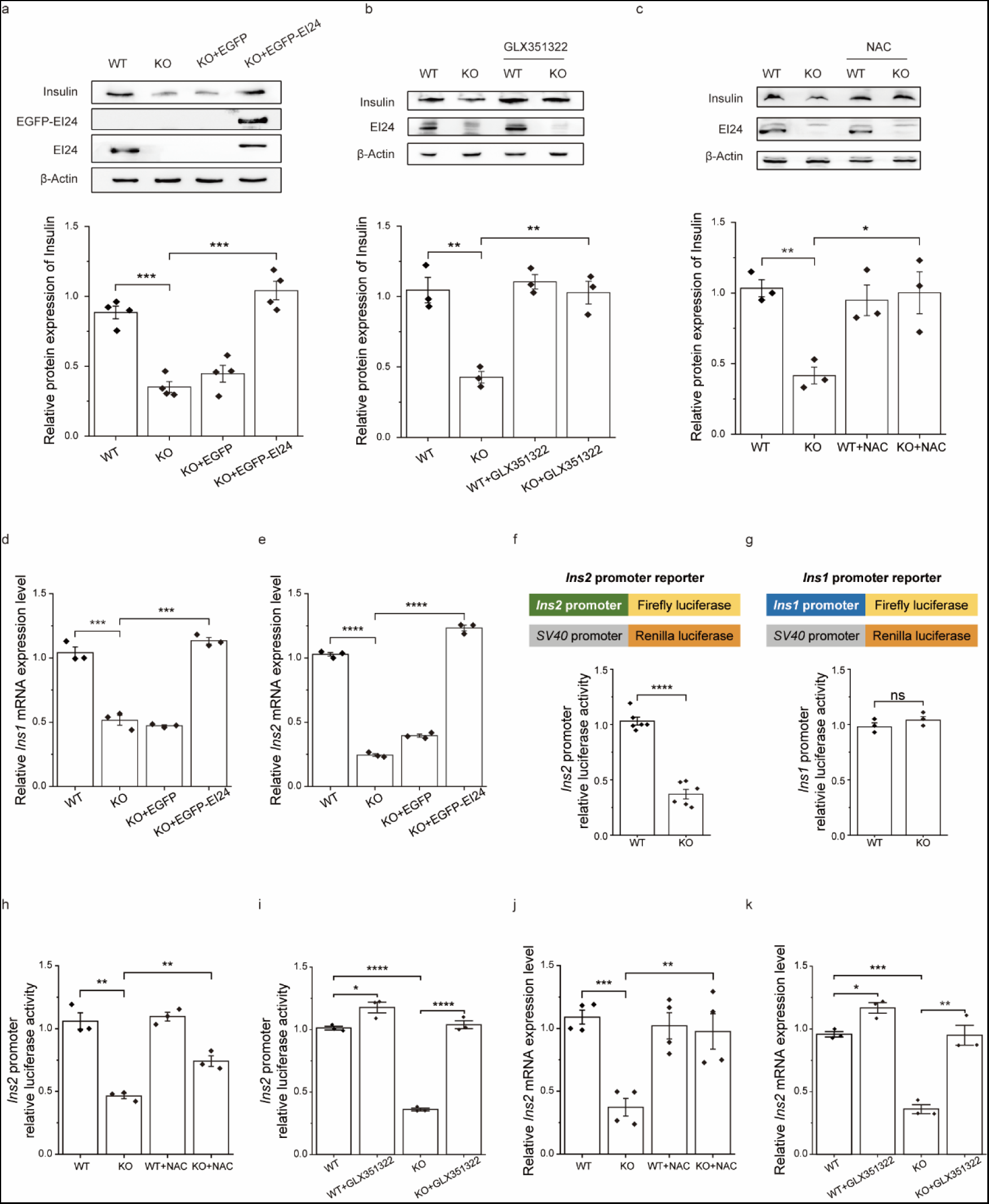
Loss of EI24 decreases insulin secretion through inhibition of its transcription. a. Levels of insulin in WT and EI24 KO INS-1 cells. b and c. Level of insulin in WT and EI24 KO INS-1 cells with or without NOX4 inhibitor (b) or NAC (c) treatment. d and e. Relative *Ins1* (d) and *Ins2* (e) mRNA levels in WT and EI24 KO INS-1 cells. f. Transcriptional activity of *Ins2* (f) and *Ins1* (g) genes using the dual fluorescence reporter system. h and i. Transcriptional activity of *Ins2* in WT and EI24 KO INS-1 cells with or without NAC (h) or NOX4 inhibitor (i) treatment. j and k. Relative *Ins2* mRNA level in WT and EI24 KO INS-1 cells with or without NAC (j) or NOX4 inhibitor (k) treatment. Data are expressed as the mean ± S.E.M. \**p* < 0.05, \*\**p* < 0.01, \*\*\**p* < 0.001, \*\*\*\**p* < 0.0001.

mRNA-seq data showed that the levels of insulin mRNA (*Ins1* and *Ins2*) were substantially reduced (Figure S6). To gain further insight into the relationship between H_2_O_2_ and insulin, we first confirmed the levels of *Ins1* and *Ins2* mRNA by quantitative PCR in WT and KO cells. Consistent with the results of mRNA-seq, deletion of EI24 produced a reduction in *Ins1* and *Ins2* mRNA to 49.44% and 23.8% of the WT controls, respectively (Figure 5d, e). Additionally, over-expression of EI24 (but not the EGFP control) rescued *Ins1* and *Ins2* mRNA levels in EI24 KO cells (Figure 5d, e). Using the dual fluorescence reporter system, we found that EI24 primarily regulates the transcription of *Ins2* mRNA (Figure 5f), but not *Ins1* (Figure 5g). Moreover, treating with NAC or a NOX4 inhibitor could boost the transcription of Ins2 mRNA (Figure 5h, i) and elevate its level to match that of WT cells (Figure 5j, k). Next, we aimed to identify key transcription factors for EI24-regulated insulin transcription by capturing proteins that bind specifically to the *Ins2* promoter. Among the proteins identified by mass spectrometry (Table S3), we observed a significant reduction of MAFA, a beta cell-specific transcription factor, in the EI24 KO group (Figure 6a, Table S3). The western blot analysis of samples obtained from the DNA affinity purification confirmed that MAFA indeed capable of binding to the *Ins2* promoter (Figure 6b). Furthermore, the amount of MAFA immunoprecipitated exhibited a notable reduction in KO cells (Figure 6b). As there was no change in the MafA mRNA levels following EI24 depletion when compared to control cells (Figure S7), we proceeded to analyze the levels of MAFA protein expression. Our findings revealed a significant decrease in MAFA within the EI24 KO group (Figure 6c). Overexpression of EI24 successfully rescued the expression level of MAFA to similar levels as the WT group, while expressing EGFP alone failed to do so (Figure 6c). The decrease in MAFA expression observed in EI24 KO cells cannot be attributed to protein degradation. This is supported by our findings that treatment with CHX, an inhibitor of protein synthesis, did not result in differences in protein degradation rates between WT and EI24 KO cells. (Figure S8a). Moreover, the level of MAFA remained unchanged in both the WT and KO groups, regardless of MG132 treatment (Figure S8b). As a result, our focus turned towards investigating the regulation of *MafA* translation by EI24. According to the data presented in the Figure 6d, the level of MAFA in WT cells showed a rapid increase, reaching half of the initial level within 2 hours after CHX removal, whereas there was no change in the MAFA level in EI24 KO cells. These findings demonstrate that EI24 is likely involved in regulating the translation of MAFA, impacting its overall abundance. Additionally, when treated with a NOX4 inhibitor, the expression of MAFA protein in the EI24 KO cells reached a level comparable to that observed in WT cells (Figure 6e). Similarly, treatment with NAC both in WT and EI24 KO INS-1 cells resulted in an increase in MAFA levels when compared with the non-treatment group (Figure 6f). To investigate the potential effects of NAC on glucose metabolism control, the mice were administered daily doses of NAC in drinking water and subsequently underwent glucose tolerance tests and insulin tolerance tests. The results showed that NAC delivery improved glucose tolerance in both male and female mice with pancreatic beta-cell-specific EI24 KO (Figure 6g, h), while it had no obvious effect on insulin sensitivity (Figure 6i, j). These findings indicate that the removal of excess ROS by NAC effectively enhances pancreatic beta-cell function in EI24 KO mice.

**Figure 6:**
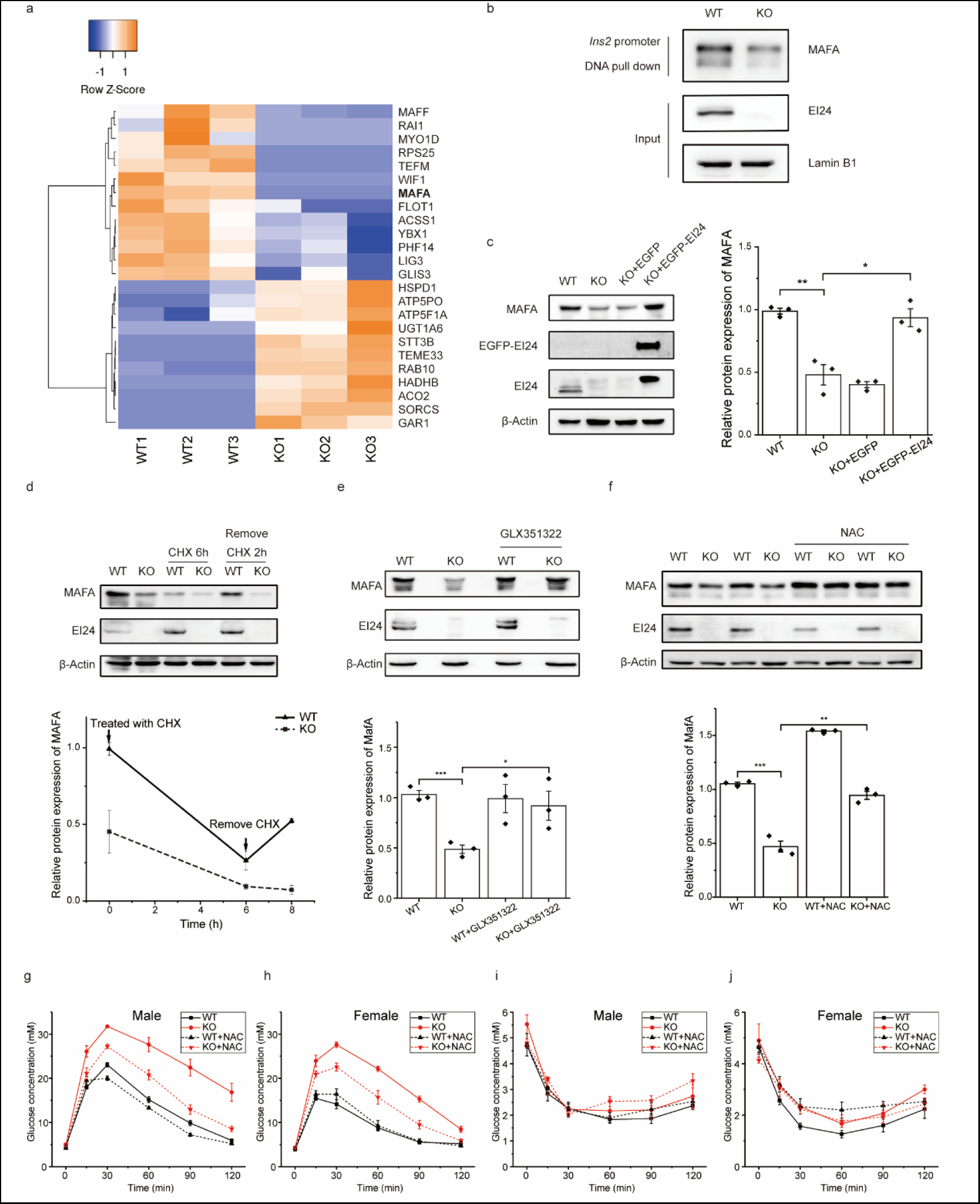
Loss of EI24 inhibits *MafA* translation by relieving *Nox4* translational inhibition. a. Heatmap of genes identified by IP using *Ins2* promoter with differential expression between EI24 KO INS-1 cells and WT group. b. *In vitro* verification that MAFA can be IPed by the *Ins2* promoter. c. Protein levels of MAFA in WT and EI24 KO INS-1 cells. d. Amount of MAFA in WT and EI24 KO INS-1 cells with or without CHX treatment. e and f. Amount of MAFA in WT and EI24 KO INS-1 cells with or without NOX4 inhibitor (e) or NAC (f) treatment. g and h. Glucose tolerance tests of male (g) and female (h) mice. i and j. Insulin tolerance tests of male (i) and female (j) mice. CHX, cycloheximide; NAC, N-acetyl cysteine. Data are expressed as the mean ± S.E.M. \**p* < 0.05, \*\**p* < 0.01, \*\*\**p* < 0.001, \*\*\*\**p* < 0.0001.

## Materials and methods

### Animals

All animal experiments were approved by and performed in accordance with the guidelines of the Animal Care and Use Committee of the Institute of Biophysics, Chinese Academy of Sciences (Approval code: SYXK2022142).

*Ei24*^flox/flox^ mice (kindly provided by Prof. Hong Zhang, Institute of Biophysics, Chinese Academy of Sciences) and Ins2-Cre knockin mice (kindly provided by Prof. Weiping Zhang, Department of Pathophysiology, Second Military Medical University) were housed in colony cages with a 12-hour light/12-hour dark cycle. Pancreatic beta-cell specific *Ei24* gene KO mice were generated and crossed as previously described(Yuan et al., 2018). For NAC treatment, 8-week-old mice were administered NAC daily at 10 mg/mL in drinking water for 4 weeks. For glucose tolerance or insulin tolerance tests, the mice were transferred to a cage with food withdrawn for 16 h or 12 h, respectively. Blood samples were collected from the tails at 0, 15, 30, 60, 90, and 120 min after administration of 20% glucose or insulin based on body weight. The glucose levels were measured using a glucometer (ACCUCHEK Active, Roche Applied Science).

### Islet separation and purification

The procedure was described in detail previously(Yuan et al., 2018). In brief, 0.5 mg/mL collagenase was perfused into the pancreas through the common bile duct. The engorged pancreas was removed carefully and digested in a 37°C water bath for 25 min, following vigorous shaking until the suspension became homogeneous. The islets were isolated, picked by hand, and incubated in Hank’s buffer (Sigma, MA, USA) at 37°C with 5% CO_2_ overnight.

### Cell culture and transfection

Primary pancreatic beta cells were dispersed from the purified islets using trypsin (GIBCO, 25200072) digestion and cultured in RPMI 1640 medium (GIBCO, 11835-030) containing 10% heat-inactivated fetal bovine serum (FBS, HyClone, Logan, UT) with 1 mM sodium pyruvate, 10 mM HEPES, 2 mM glutamine, and 50 μM β-Mercaptoethanol (GIBCO, 21985023) in an incubator at 37°C with 5% CO_2_.

INS-1 cells were cultured in RPMI 1640 Medium (GIBCO) supplemented with 10% FBS, 1 mM sodium pyruvate solution, 50 μM β-Mercaptoethanol (GIBCO), and 0.1% Mycoplasma prevention reagent (Transgen, Beijing, China) at 37°C and 5% CO_2_.

Plasmid transfections were performed using Lipofectamine 3000 (Thermo Fisher Scientific, no. L3000015) for INS-1 cell lines, following the manufacturer’s instructions. RTRAF siRNAs were purchased from RiboBio (Guangzhou, China) and transfected into INS-1 cells using Lipofectamine RNAiMAX Reagent (Thermo Fisher Scientific, no. 13778150). Unless otherwise noted, cells were collected 24–36 h post-transfection.

### Plasmid constructs

The EGFP-EI24 plasmid was constructed as previously reported(Yuan et al., 2018). For the construction of pcDNA3.1-HyPer_cyto_, the open-reading frames (ORFs) of HyPer were obtained from the pLVX-HyPer plasmid (gift from Prof. Xianen Zhang, Institute of Biophysics, Chinese Academy of Sciences). The sequence for HyPer in frame with the nuclear export signal was then inserted into the NheI and NotI digested backbone of pcDNA3.1. The KDEL peptide sequence in frame and immediately downstream of the HyPer was inserted to obtain pcDNA3.1-HyPer_ER_. pcDNA3.1-Grx1-roGFP2_cyto_ and pcDNA3.1-rosfGFPiE_ER_ were gifted by Prof. Chang Chen (Institute of Biophysics, Chinese Academy of Sciences) and Prof. Lei Wang (Institute of Biophysics, Chinese Academy of Sciences), respectively. The sequence of MS2 was synthesized, amplified, and inserted into the BamHI/XbaI sites of pcDNA3.1 to generate pcDNA3.1-MS2, and the sequence of MCP-NLS was synthesized, amplified, and inserted into the XhoI/KpnI sites of pEGFP-C1 To construct EGFP-MCP-NLS plasmid. The ORF sequence of *Rtraf* was synthesized (Tsingke Biotechnology Co., Ltd.), amplified, and cloned into the XhoI/EcoRI sites of pEGFP-N1 to generate pEGFP-RTRAF. Sequences encoding the *MafA* transcript were amplified from the cDNA of INS-1 cells and cloned into the XhoI/NotI sites of pFLAG-c1 vector to generate pFLAG-MafA plasmids. pGL6-Ins1-promoter-Luc was created by inserting the sequences of *Ins1* promoter into the XhoI/MluI sites of pGL6-Luc (Beyotime Biotechnology). To generate pGL6-Ins2-promoter-Luc, *Ins1* promoter in the pGL6-Ins1-promoter-Luc plasmid was replaced with *Ins2* promoter. To examine the UTR regions of NOX4 and their impact on translation, the plasmid psiCHECK2, a gift from Prof. Xiaorong Zhang (Institute of Biophysics, Chinese Academy of Sciences), containing renilla luciferase and firefly luciferase was used. For the construction of ^5’UTR^NOX4-psiCHECK2, the synthesized *Nox4* 5’-UTR sequence was inserted into the NheI digested backbone of psiCHECK2 in frame. The synthesized *Nox4* 3’UTR sequence was inserted into the XhoI/NotI sites of psiCHECK2 to generate ^3’UTR^NOX4-psiCHECK2. Restriction enzymes were purchased from Thermo Fisher Scientific. The targeting of the indicated subcellular compartment by the plasmids was consistent with the pattern of fluorescent signal obtained from living cells.

### Flow cytometry

The day after culturing, pancreatic beta cells (about 50–60 islets) were dispersed and incubated for an additional 1 h under the corresponding treatment conditions. After that, 5 μM of CellROX™ Green (0.5 μM for INS-I cells; Thermo Fisher Scientific) was added for 45 min at 37°C. The cells were washed twice with PBS, and 5 μM of SYTOX™ Red was added to label dead cells. CellROX™ fluorescence intensity was measured on an FACSAria IIIu flow cytometer (BD Biosciences) using 488 nm and 640 nm lasers. FlowJo software (BD Biosciences) was utilized to analyze flow cytometry data.

To analyze the level of H_2_O_2_ or thiol redox, INS-1 cells were transfected with HyPer or roGFP2 expression vectors using Lipofectamine 3000 (Thermo Fisher Scientific) according to the manufacturer’s instructions. Flow cytometry experiments were performed 24–36 h after transfection. During flow cytometry analysis of the samples, the cells were gated, and 488 to 405 nm signal ratios for Hyper as well as 405 to 488 nm signal ratios for roGFP2 were determined, respectively.

### ROS measurement

INS-1 cells were stained with 10 µM fluorescent dye 2′,7′-dichlorodihydrofluorescein diacetate (DCFH-DA) in Hank’s buffer for 20 min at 37°C following treatment. The cells were then washed three times with PBS and imaged using an FV1200 Laser Scanning Confocal Microscope (Olympus) with 488-nm lasers.

### Immunostaining assays

Cells seeded on fibronectin-coated glass cover slips were fixed with 4% PFA for 15 min at 37°C and permeabilized in 0.1% Triton X-100 (Sigma) for 5 min at room temperature. After blocking with 5% goat serum for 1 h at room temperature, the cells were incubated with indicated first antibodies, followed by washing with PBS for three times and incubation with fluorescence-labeled secondary antibodies. Images were acquired using an Olympus FV1200 Laser Scanning Confocal Microscope (Olympus) with a 60× (NA = 1.40) oil objective. The images were quantified and analyzed using the ImageJ software (National Institutes of Health).

### mRNA sequencing

Multiplex analysis of Poly(A)-linked sequences (MAPS) was carried out as previously described with minor modifications(Zhou et al., 2014). In brief, total RNA (1 μg) from islets was added to 50 μM first-strand RT primer (the sequence at the 5’ portion corresponds to primer P7 on the Illumina flowcell), 10 mM dNTP mix, and distilled water to a total of 10 μL. The mixture was then heated for 5 min at 65°C and subsequently incubated on ice for at least 1 min. After the samples were cooled, 10× RT buffer, 25 mM MgCl_2_, 0.1 M DTT, 40 U/μL RNase inhibitor, and 200 U/μL SuperScript III (Invitrogen) were added to the mixture. The mixture was incubated at 50°C for 50 min and terminated at 85°C for 5 min. Then, the first-strand cDNA was purified with the NucleoSpin gel and PCR clean-up kit (Macherey-Nagel, no. 470609) following the manufacturer’s instruction. The purified cDNA was added to a mixture of 10× terminal transferase buffer (NEB, no. M0315S), 2.5 mM CoCl_2_, 10 mM ddNTP, and 20 U/μL terminal transferase (NEB, no. M0315S). After incubation at 37°C for 30 min and termination at 70°C for 10 min, the mixture was incubated with washed magnetic beads (Invitrogen, no. 65001) for 30 min, and then, the beads were collected and washed with 0.1 M NaOH, followed by incubation for 5 min at room temperature. The second-strand cDNA was synthesized by adding 100 μM primer (the specific sequence at the 5′ portion corresponds to the primer for sequencing on the Illumina flowcell), 10× Taq DNA Polymerase Buffer, 10 mM dNTP mix, and 5 U/μL Taq DNA polymerase (NEB) for incubation at 25°C for 60 min and heating at 68°C for 30 s and then at 75°C for 5 min. After washing with a buffer, the tubes containing beads were heated at 95°C for 5 min and immediately transferred to a magnetic stand to collect the supernatant containing the released second-strand cDNA. Next, PCR was performed, and the PCR products were purified for multiplex sequencing. Data were generated by MAPS by using the software package maps3end.

### DNA promoter pull-down assay

To determine the transcription factors associated with the *Ins2* promoter, a standard procedure was used as previously described with slightly modification(Chaparian and van Kessel, 2021). Briefly, cell nuclei were isolated, diluted in buffer D (20 mM HEPES, pH 7.9, 100 mM KCl, 0.2 mM EDTA, 8% glycerol with inhibitor cocktail), aliquoted, and stored at −80°C. Using pGL6/*Ins2-*promoter/Luc as the template, the DNA probe was prepared by PCR using 5′ biotinylated primers and was purified using a gel extraction kit (Cwbiotech, JiangSu, China). After washing three times with 2×B/W buffer (20 mM HEPES pH 7.5, 10 mM CaCl_2_, 100 mM KCl, 25% glycerol), Dynabeads M-280 Streptavidin (Invitrogen, 11206D) were incubated with DNA probe (∼15 µg DNA) for 40 min at room temperature. The bead-DNA complex was washed with 0.5 M TE buffer (pH 8.0) three times and then with BS/THES buffer (50 mM Tri-HCl, pH 7.5, 10 mM EDTA, 20% Sucrose, and 140 mM NaCl) three times. The bead-DNA complex was gently mixed with nuclear lysate at 4°C for 80 min. The beads were collected using a magnetic stand and washed with BS/THES buffer five times. Elution buffer (0.1% SDS) was added and incubated at 95°C for 5 min. The supernatant sample was collected and prepared for SDS-PAGE and mass spectrometric analysis.

### Dual-luciferase report system

To identify the transcriptional level of *Ins1* or *Ins2*, INS-1 cells were co-transfected with the pGL6-Ins1-promoter-Luc reporter plasmid and pRL-SV40-C plasmid as control or the pGL6-Ins2-promoter-Luc reporter plasmid and pRL-SV40-C plasmid, respectively. After 24–36 h, the samples were prepared and analysis was performed using the Dual-Lumi™ reporter gene assay kit (Beyotime Biotechnology) following the manufacturer’s instructions.

In the system assessing the relative reporter activities of *Nox4* mRNA’s different UTRs, the firefly luciferase signal serves as an expression normalization control, while the renilla luciferase signal serves as the reporter in a single-read assay.

### Sample preparation for quantitative proteomic analysis

Equal amounts of protein from WT and EI24 KO INS-1 cells were separated by SDS-PAGE. The bands were excised, reduced with 25 mM of DTT, and alkylated with 55 mM iodoacetamide, followed by in-gel digestion with sequencing-grade modified trypsin overnight at 37°C. The peptides were extracted twice with 0.1% trifluoroacetic acid in 50% acetonitrile aqueous solution for 30 min and dried in a speedvac. After being redissolved in 25 μL 0.1% trifluoroacetic acid, 6 μL of the extracted peptides was analyzed by Q Exactive HF-X mass spectrometer.

### Mass spectrometric analysis

The peptides were separated by a 120-min gradient elution at a flow rate of 0.30 µL/min with a Thermo-Dionex Ultimate 3000 HPLC system, which was directly interfaced with Q Exactive HF-X mass spectrometer (Thermo Fisher Scientific, Bremen, Germany). The analytical capillary column was fused silica (75 µm ID, 150 mm length; Upchurch, Oak Harbor, WA) and packed with C-18 resin (300 Å, 5 µm, Varian, Lexington, MA). The mobile phases A and B consisted of 0.1% formic acid and 100% acetonitrile with 0.1% formic acid, respectively. The Q Exactive HF-X mass spectrometer was operated in a data-dependent acquisition mode using Xcalibur 4.5 software. There was a single full-scan mass spectrum in the orbitrap (350–1500 m/z, 60,000 resolution), followed by 2 seconds data-dependent MS/MS scans in an ion routing multipole at 30% normalized collision energy. The MS/MS spectra from the LC-MS/MS runs were searched against the Rat UniProt using an in-house Proteome Discoverer (Version PD1.4, Thermo-Fisher Scientific, USA).

### Immunoprecipitation

INS-1 cells were transfected with the appropriate plasmids. After 24 h, the cells were collected and lysed on ice in RIPA buffer containing protease inhibitor cocktail (Sigma) and then centrifuged at 12,000 rpm for 5 min at 4°C. The supernatants were incubated with GFP-Trap_A beads (ChromoTek, Thermo Fisher Scientific) on a rotator for 2 h at 4°C. The precipitated samples were washed three times with washing buffer (50 mM Tris–HCl, 300 mM NaCl, 1 mM EDTA, 1% Triton X-100, pH 7.4) and eluted in 0.1 M glycine (pH 2.0). After neutralization by 1 M Tris base, the immunoprecipitated samples were utilized for the following experiments. For the RNA IP, the 3’-UTR domain of NOX4 fused with MS2 and EGFP-labeled MCP was co-transfected into INS-1 cells. As a result of the interaction between MS2 and MCP, the proteins interacting with the 3’-UTRs of NOX4 were pulled down and analyzed using specific antibodies. Alternatively, the RNA species that were immunoprecipitated by the EGFP-labeled protein of interest were purified as previously reported(Yoon et al., 2012)and analyzed by PCR.

### Western blotting

Whole-cell lysates prepared using cold RIPA buffer (Beyotime Biotechnology, Shanghai, China) containing a proteinase inhibitor mixture (Sigma) were quantitated using the Bradford assay (Bio-Rad). The protein samples were separated by SDS-PAGE and transferred onto PVDF membranes (Millipore, Billerica, MA, USA). The membranes were incubated with primary antibodies against EI24 (Sigma), Insulin (Cell Signaling Technology), MAFA (Cell Signaling Technology), NOX4 (Novus Biologicals), ERO1α (R&D Systems), PRDX4 (R&D Systems), GPX7 (ABclonal), GPX8 (Novus Biologicals), RTRAF (Proteintech), Lamin B1 (Proteintech), β-tubulin (Proteintech), and β-actin (Cell Signaling Technology), followed by incubation with the appropriate HRP conjugated secondary antibodies (Sungene Biotech, Tianjin, China). Protein expression was detected using enhanced luminescence reagents (Thermo Scientific, Waltham, MA, USA).

## Discussion

It’s worth noting that the expression of NOX enzymes is tightly regulated in early life. Dysregulation of NOX enzyme expression may lead to an increase in ROS and the onset of disease(Lambeth, 2007). Understanding the regulatory mechanisms that control the expression of NOX enzymes, especially the negative mechanisms that inhibit their expression, can help uncover the development of diseases such as diabetes. The ability of NOX4 to constitutively produce H_2_O_2_ makes it unique among NOX family members(Wang et al., 2023). Current research into the molecular mechanisms that control NOX4 expression is focused primarily at the transcriptional level(Hecker et al., 2023), and there are few studies on translation. HuR is the only reported positive regulator upregulating NOX4 expression in high glucose-stimulated beta cells(Shi et al., 2020). However, it is not known how NOX4 is regulated at the translation level when cells are in the basal state. Our study presents compelling evidence that EI24 interacts with the RNA-binding protein RTRAF, recruiting it to the 3’-UTR of *Nox4* mRNA and thereby inhibiting the translation of *Nox4*. This process is crucial for maintaining a low level of basal H_2_O_2_. To our knowledge, EI24 is the first protein identified that negatively regulates *Nox4* translation. Interestingly, the expression level of EI24 changed in response to the concentration of H_2_O_2_, suggesting that EI24 may also play a role in maintaining H_2_O_2_ homeostasis when cells are under oxidative stress.

RTRAF (aka hCLE/C14orf166) is an RNA-binding protein with both nuclear and cytoplasmic localization(Huarte et al., 2001), which modulates RNA synthesis and processing. Research has demonstrated that RTRAF is part of protein complexes that shuttle between the nucleus and cytoplasm and plays a role in transporting RNA molecules between these two cellular compartments(Pérez-González et al., 2014). Furthermore, the hCLE/RTRAF-HSPC117-DDX1-FAM98B complex exhibits cap-binding activity and enhances mRNA translation(Pazo et al., 2019). In the current study, we found that knockdown of *Rtraf* results in increased translation of *Nox4* mRNA in WT INS-1 cells. Deletion of EI24 leads to the relocation of RTRAF from the cytoplasm to the nucleus, disrupting its interaction with the 3′UTR of *Nox4* mRNA and ultimately releasing the translational repression of *Nox4* mRNA. Hence, our research reveals a new role of RTRAF in the negative regulation of *Nox4* mRNA translation, and this inhibitory function of RTRAF in translation may be conserved in other tissues. In developing brain, RTRAF has been found to be the core element of cytosolic, ribosome-containing RNA granules that transport specific mRNAs from the cell body to the dendrites, allowing local mRNA translation at sites distant from the nucleus(Elvira et al., 2006). It is currently unclear how the translation of RTRAF-transported mRNA is inhibited before reaching its destination. Although not yet confirmed, our work may offer a potential explanation. RTRAF might have a similar role in inhibiting translation during RNA transport, while EI24 and/or other factors could be involved in regulating this through interaction with RTRAF.

Maintaining a delicate equilibrium between the production of H_2_O_2_ and the antioxidant defense is crucial for the health and insulin production of pancreatic beta cells. H_2_O_2_ increases the level of the homeodomain transcription factor PDX1 (pancreatic and duodenal homeobox 1)(Baumel-Alterzon and Scott, 2022), a key transcription factor necessary for beta-cell development and the expression beta-cell specific genes. Additionally, H_2_O_2_ plays a crucial role in glucose stimulated insulin secretion(Plecita-Hlavata et al., 2020). However, under pathological conditions like chronic hyperglycemia and glucolipotoxicity, the rise in ROS overwhelms cellular defenses, leading to permanent changes in biomolecules and disrupting proteostasis. This ultimately adds to cellular dysfunction and damage. The translational repressive regulation of *Nox4* by EI24 through interacting with RTRAF, as discovered in the current study, is crucial in maintaining a low level of H_2_O_2_ for the normal function of pancreatic beta cells. Additionally, the interaction between EI24 and RTRAF may also play a protective role when beta cells are under pathological conditions, such as high glucose stimulation. It has been reported that under high glucose stimulation, HuR upregulates *Nox4* translation, resulting in increased H_2_O_2_ production(Shi et al., 2020).

This suggests that in beta cells stimulated by high glucose, excessive nutrient availability may lead to increased synthesis and subsequent higher levels of H_2_O_2_ in the ER for oxidative protein folding, especially in cells with strong secretion(Appenzeller-Herzog, 2011; Lenzen, 2017). This mechanism could offer an explanation for the impairment of pancreatic beta cells in patients with type 2 diabetes under pathological conditions. We hypothesize that the reduced expression of EI24 in diabetic animal models(Yuan et al., 2018) or patients may disturb the balance of ROS and result in an increase in H_2_O_2_, ultimately leading to impaired pancreatic beta cell function, reduced insulin synthesis, and secretion.

In pancreatic beta cells, several core transcription factors, including PDX1, Neurogenin-3, and MAFA, respond to various physiological stimuli to regulate the insulin gene’s transcription(Zhu et al., 2017). However, the exact mechanisms by which ROS affect these transcription factors and their impact on insulin transcription are still largely unknown. In our current study, we observed an interaction between the *Ins2* promoter and MAFA, PDX1, and other transcription factors in INS-1 cells. Interestingly, after EI24 knockout, only MAFA dissociated from the insulin promoter. As EI24 knockout leads to an increase in intracellular H_2_O_2_ (Figure 1f), and the addition of a NOX4 inhibitor or the non-specific reducing agent NAC can restore the transcription and synthesis of insulin (Figure 5b, c), we suspect that the key transcription factors may employ different mechanisms to regulate the transcription and synthesis of insulin in response to oxidative stimulation. We also propose that EI24-mediated H_2_O_2_ stress regulates insulin gene transcription by influencing the interaction between MAFA and the *Ins2* promoter.

In conclusion, we have uncovered a new mechanism in which EI24 serves as a negative regulator, controlling the translation of *Nox4* and influencing the localization of RTRAF and its interaction with *Nox4* mRNA. This mechanism plays a crucial role in maintaining H_2_O_2_ balance and insulin synthesis in pancreatic beta cells (Figure 7). Given the similarities between nerve cells and pancreatic beta cells in terms of peptide synthesis and secretion, as well as their intolerance to ROS due to natural catalase deficiency leading to reduced cytosolic antioxidant capacity(Benakova et al., 2021; Konno et al., 2021; Wang and Wang, 2017), we hypothesize that the translational inhibition of *Nox4* expression by EI24-RTRAF interaction may also be present in neurons. Further investigation is warranted to verify the connection between EI24- regulated NOX4 expression at the translational level and the secretory function of cells in other secretory cell types.

**Figure 7:**
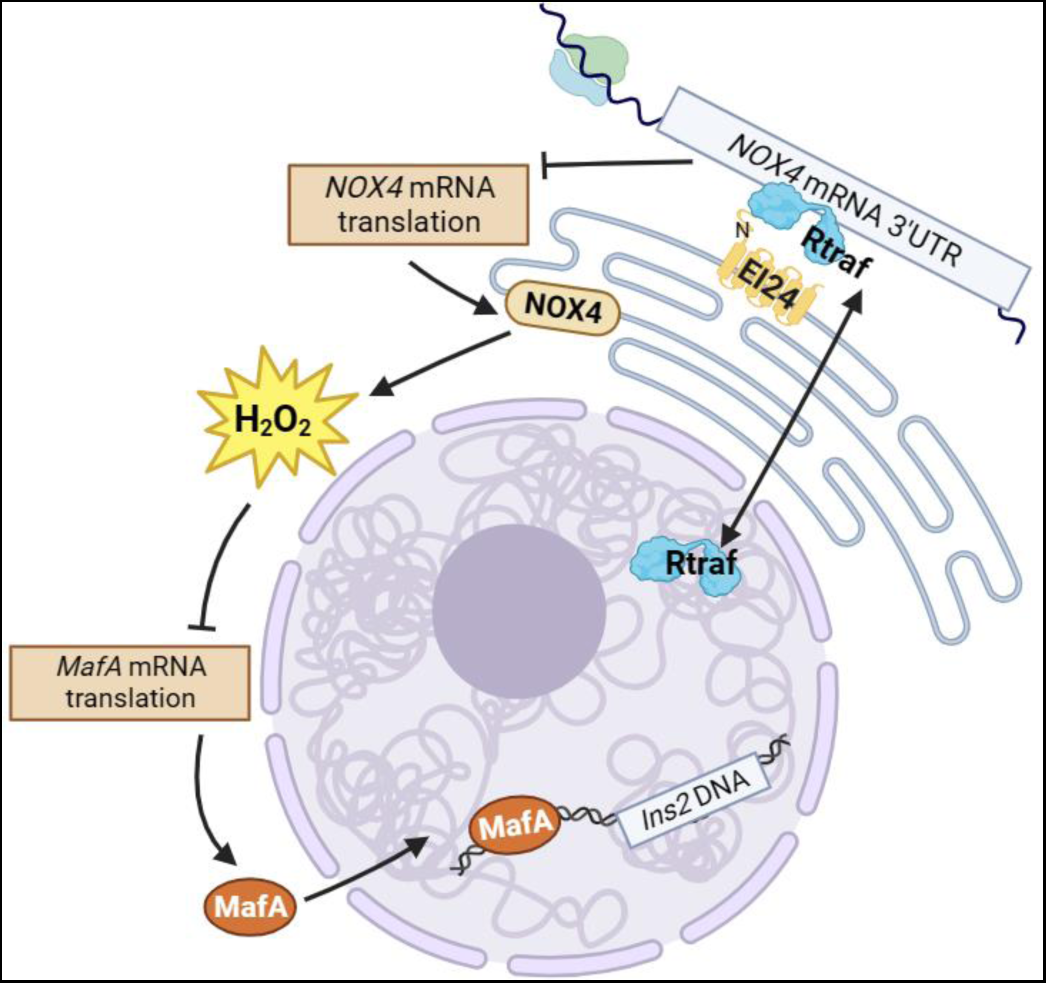
A schematic diagram illustrates the mechanism by which EI24 interacts with the RNA-binding protein RTRAF and *Nox4* mRNA 3’-UTR to inhibit the translation of *Nox4*. This inhibition is essential for maintaining basal H_2_O_2_ homeostasis and insulin synthesis

## Acknowledgements

This project was supported by the National Key R&D Program of China (2022YFC3400600 and 2022ZD0211900), the National Natural Science Foundation of China (92254306, 21927813, 32227802, 31970704 and T2394513), the Strategic Priority Research Program of Chinese Academy of Sciences (XDB37040301). We thank Dr. Hong Zhang (Institute of Biophysics, Chinese Academy of Sciences) and Dr. Weiping Zhang (Department of Pathophysiology, Second Military Medical University) for the mice of *Ei24*^flox/flox^ and Ins2-Cre, respectively. We thank Dr. Lei Wang (Institute of Biophysics, Chinese Academy of Sciences) for plasmid pHyPer-ER, Dr. Xianen Zhang (Institute of Biophysics, Chinese Academy of Sciences) for plamid pLVX-HyPer, Dr. Chang Chen (Institute of Biophysics, Chinese Academy of Sciences) for plasmids pcDNA3.1-Grx1-roGFP2cyto and pcDNA3.1-rosfGFPiEER, and Dr. Xiaorong Zhang (Institute of Biophysics, Chinese Academy of Sciences) for plasmid psiCHECK2. We thank Prof. Haiteng Deng and Dr. Chongchong Zhao in Proteinomics Facility at Technology Center for Protein Sciences, Tsinghua University, for protein MS analysis. We thank Dr. Hong Zhang and Dr. Xiaorong Zhang for insightful discussion.

---

**Figure S1:**
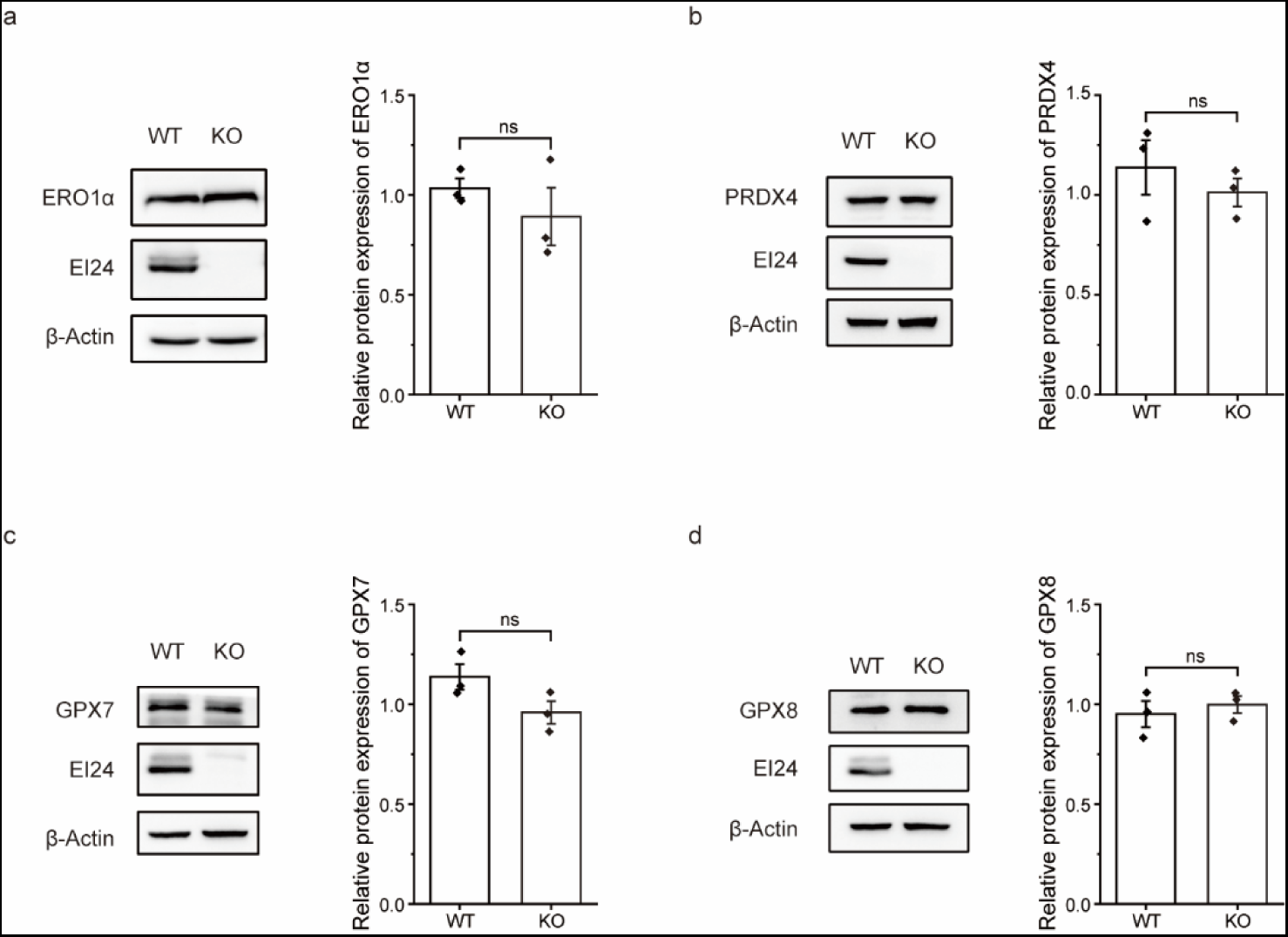
EI24 deficiency has no effect on ERO1 and antioxidant enzyme levels. The protein expression levels of ERO1α (a), PRDX4 (b), GPX7 (c), and GPX8 (d) were analyzed by Western blot. The data are presented as the mean ± S.E.M.

**Figure S2:**
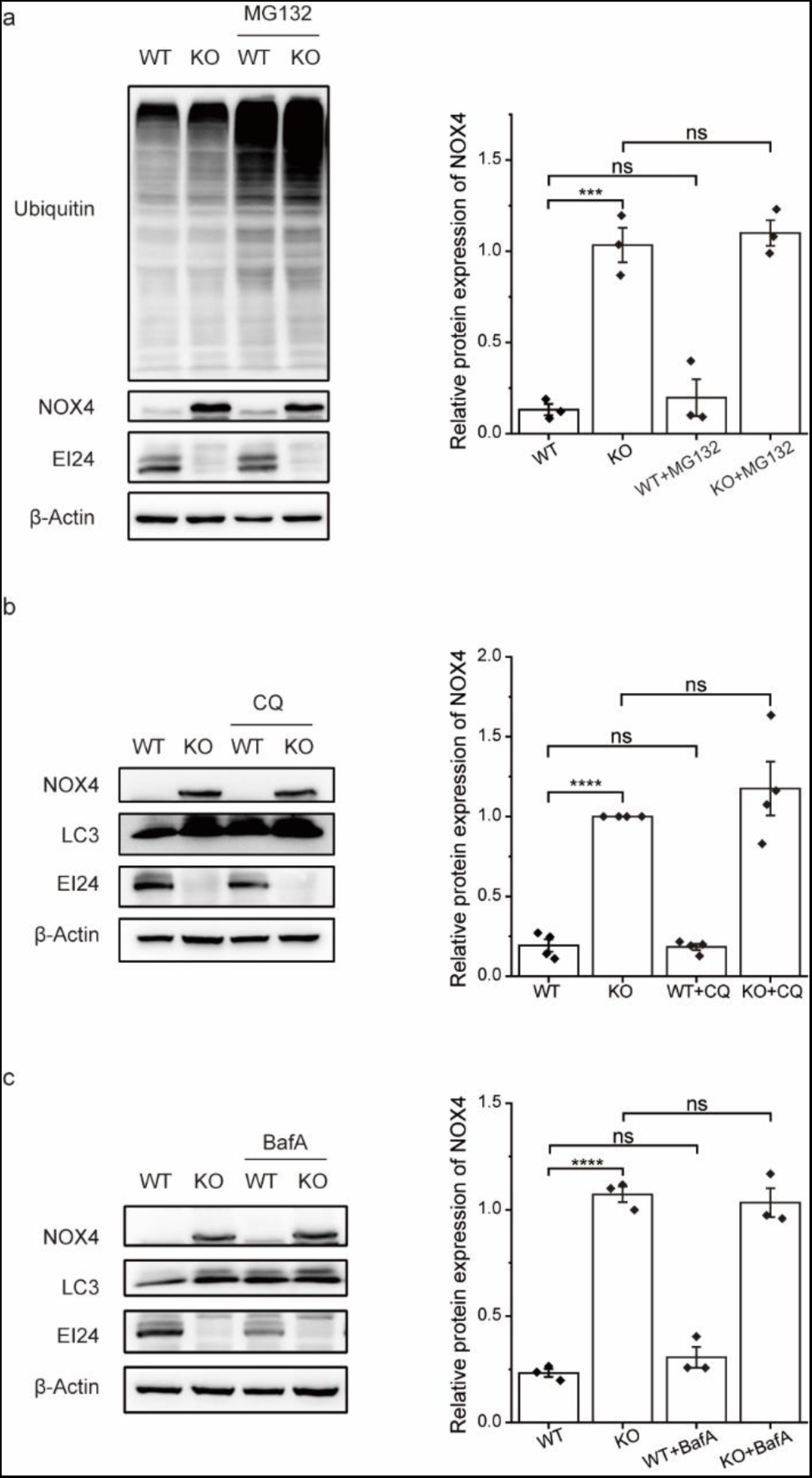
EI24 deficiency has no effect on NOX4 degradation. INS-1 cells were treated with or without MG132 (a), chloroquine (CQ, b), or bafilomycin (BafA, c). The protein expression levels were analyzed by Western blot. The data are presented as the mean ± S.E.M. \*\*\**p* < 0.001, \*\*\*\**p* < 0.0001.

**Figure S3:**
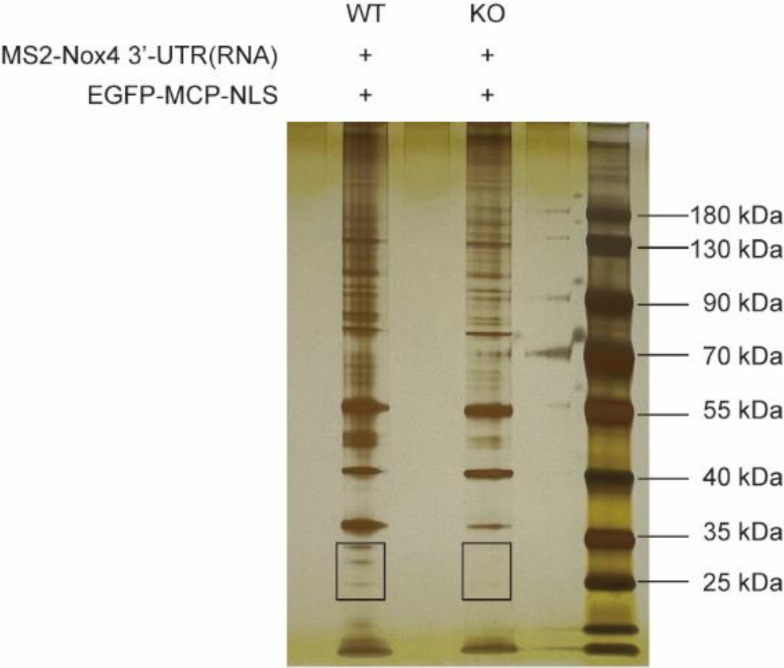
Silver staining of proteins that were immunoprecipitated using the 3’-UTR of *Nox4*. Rectangles indicate the bands that were excised for mass spectrometry analysis.

**Figure S4:**
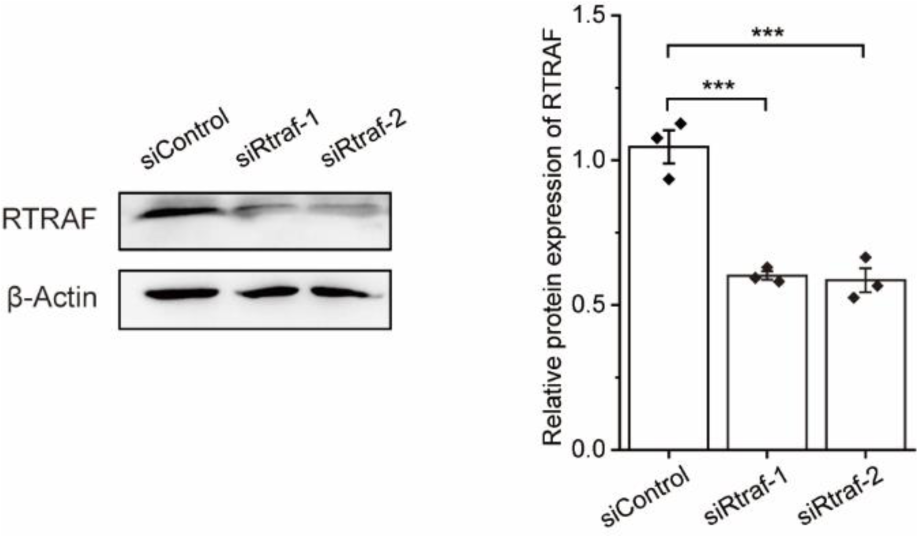
Western blot analysis of the effect of RTRAF siRNA interference on RTRAF protein expression.

**Figure S5:**
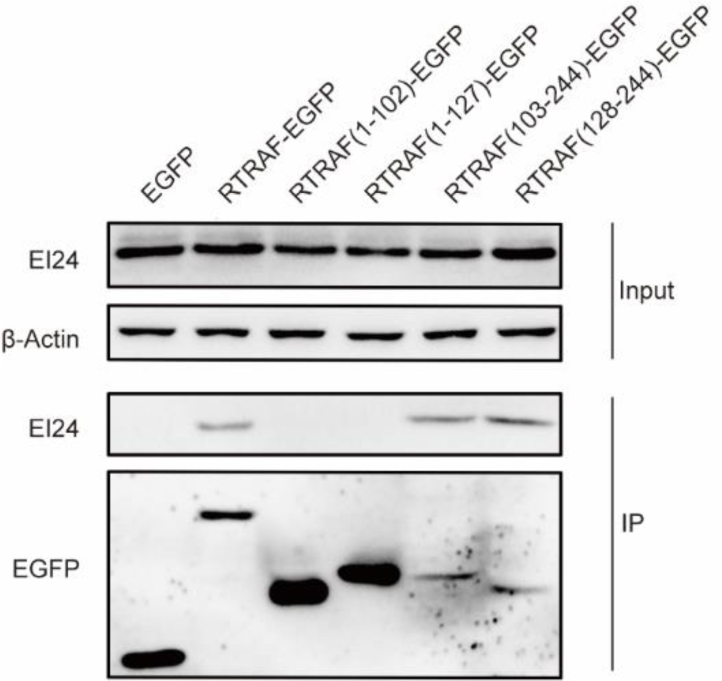
Identification of the interacting domains of EI24 and RTRAF by co-IP of EI24 using RTRAF truncation mutants.

**Figure S6:**
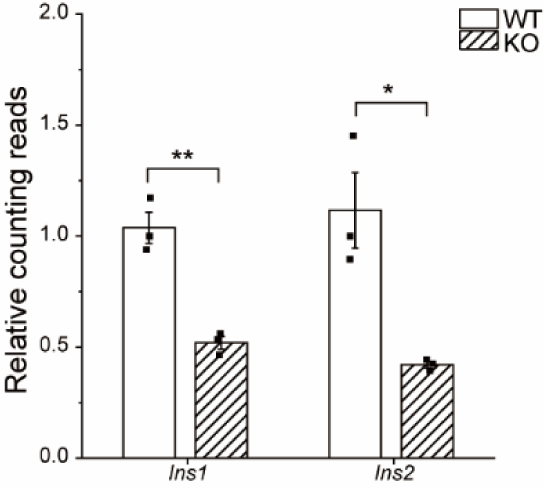
Relative counting reads of *Ins1* and *Ins2* mRNA analyzed by mRNA sequencing. The data are presented as the mean ± S.E.M. \**p* < 0.05, \*\**p* < 0.01.

**Figure S7:**
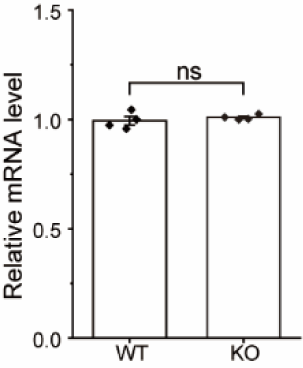
Relative *MafA* mRNA levels in WT and EI24 KO INS-1 cells. Data are expressed as the mean ± S.E.M.

**Figure S8:**
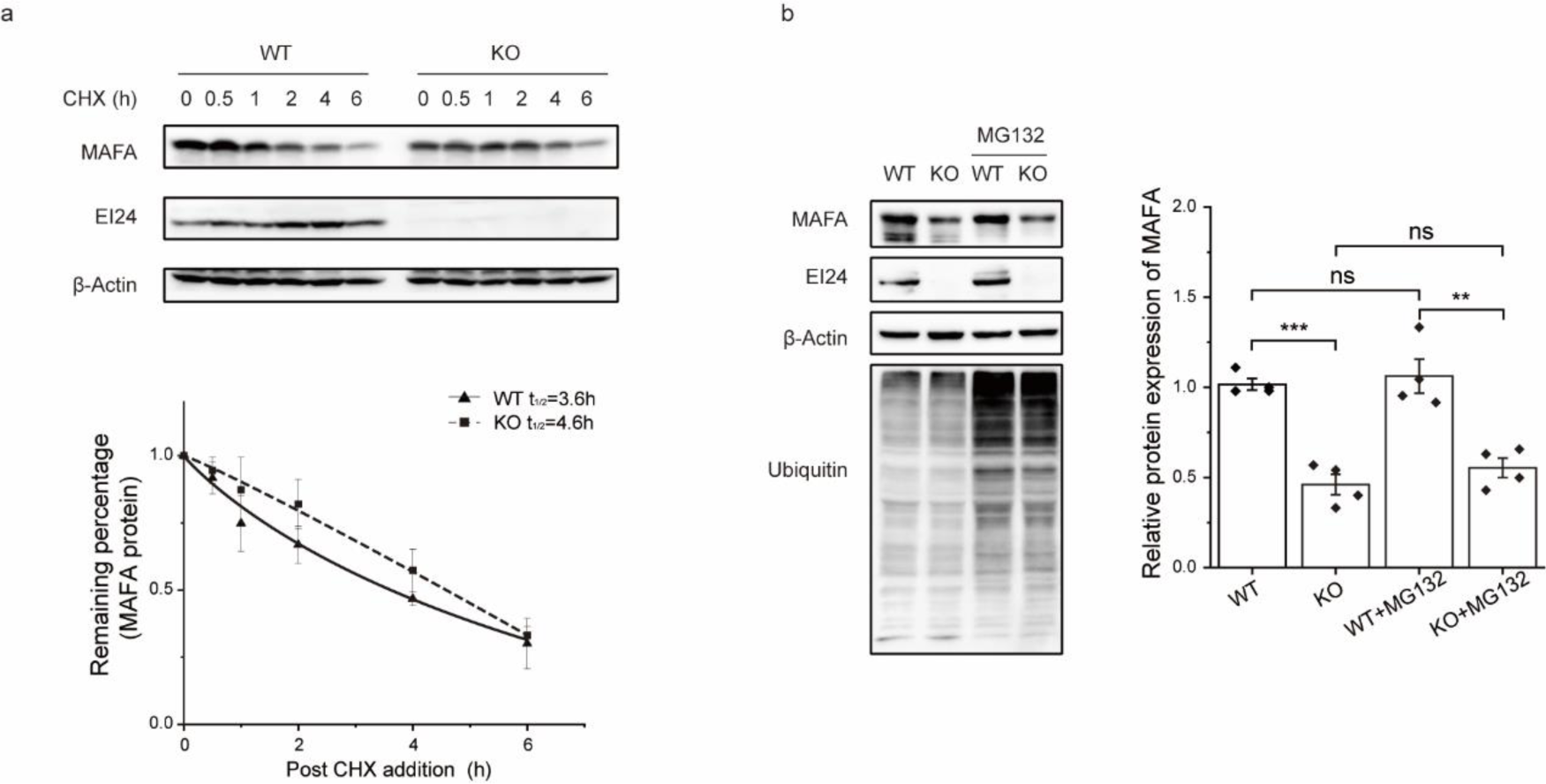
EI24 deficiency has no effect on MAFA degradation. INS-1 cells were treated with or without cycloheximide (a), or MG132 (b). Protein expression levels were analyzed by Western blot. Data are expressed as the mean ± S.E.M. \*\*\**p* < 0.001, \*\*\*\**p* < 0.0001.

